# Integrating ensemble systems biology feature selection and bimodal deep neural network for breast cancer prognosis prediction

**DOI:** 10.1101/810176

**Authors:** Li-Hsin Cheng, Che Lin

## Abstract

**Motivation:** Breast cancer is a heterogeneous disease. In order to guide proper treatment decisions for each individual patient, there is an urgent need for robust prognostic biomarkers that allow reliable prognosis prediction. Gene feature selection on microarray data is an approach to systematically discover potential biomarkers. However, common pure-statistical feature selection approaches often fail to incorporate prior biological knowledge and thus tend to select genes that lack biological insights. In addition, due to the high dimensionality and low sample size properties of microarray data, selecting robust gene features is an intrinsically challenging problem. We therefore combined systems biology feature selection with ensemble learning in this study, aiming to address the above challenges and select genes with biological insights, as well as robust prognostic predictive power. Moreover, in order to capture the complex molecular processes of breast cancer, where multiple disease-contributing genes may exist and interact, we adopted a multi-gene approach to predict the prognosis status using machine learning classifiers.

**Results:** We systematically evaluated three different ensemble approaches that all improved the original systems biology feature selector. We found that compared to the most popular data-perturbation approach, function perturbation can produce significant improvement with just a few ensembles. Among all, the hybrid ensemble approach led to the most robust feature selection result, and the identified genes were shown to be highly involved in pathways, such as ubiquitination and cell cycle. Final prognosis prediction models were constructed using the identified genes and clinical information as input features. Among all models, bimodal deep neural network (DNN) achieved the highest AUC (area under receiver operating characteristic curve) in test performance evaluation, where subsequent survival analysis also verified its ability to differentiate patients with different prognosis statuses. In summary, the study demonstrated the potential of ensemble learning to improve gene feature selection robustness, as well as the potential of bimodal DNN in providing reliable prognosis prediction and guiding precision medicine.

## 1 Introduction

Breast cancer is a heterogeneous group of tumors with variable morphologies, molecular profiles, and clinical outcomes (Polyak, 2011). Reliable prognosis prediction is thus challenging, yet essential, for a precise and personalized treatment decision. During the past decades, breast cancer biomarkers have been identified to estimate diverse responses in prognosis and therapeutic efficacy for different patients. For example, ER, PR, HER2, Ki67, and uPA/PAI-1 are some of the well-known breast cancer biomarkers that provide prognostic insights (Duffy *et al*., 2017). Joint evaluation of the immunohistochemical staining (IHC) statuses of ER, PR, and HER2 can further divide patients into subtypes, such as hormone receptor positive breast cancer (ER+/PR+) (Dunnwald *et al*., 2007) or triple negative breast cancer (ER-/PR-/HER-) (Lehmann *et al*., 2011; Carey *et al*., 2007; Dent *et al*., 2007), which are relevant for prognosis.

In order to discover more potential biomarkers to aid in reliable prognosis prediction, it is necessary to systematically analyze all possible gene candidates, which can be viewed as a feature selection problem performed on high-throughput microarray gene expression data. However, feature selection based on a pure statistical approach often fails to incorporate prior biological knowledge, and thus, tends to select genes that lack biological insights. In addition, most feature selection methods are supervised approaches which rely on labeled samples that are generally scarce. Therefore, we adopted the unsupervised systems biology feature selector (Lai *et al*., 2019) as our core feature selector. The systems biology feature selector selects genes through interaction network analysis, and two aspects of prior biological knowledge are incorporated — prognostic-relevant split criteria and BioGrid gene/protein interaction repository (Stark *et al*., 2006). The selector divides samples into two groups based on prognostic-relevant split criteria instead of classification label and constructs a gene interaction network for each group based on BioGrid. Difference analysis of two networks was carried out successively, with an output score for each gene summarizing how differently the gene interacts with its partners in two distinct prognosis statuses. The score is then used to rank and select the genes. Therefore, the gene feature selection result will serve as an extension to the previous breast cancer studies, which are inputted into the selector in the form of prognostic-relevant split criteria. Furthermore, since the selector is based on interaction network analysis, its feature selection result would help in understanding the molecular mechanisms of breast cancer from a topological aspect.

Another challenge of gene feature selection arises from the properties of microarray data. Usually, microarray datasets come with extremely high dimension but low sample size. The feature selection result obtained under this circumstance is often unstable, which would be highly sensitive to the given data and can thus fail to provide equally good predictive performance on unseen samples (Kalousis *et al*., 2007; Kim, 2009). To alleviate the problem of instability caused by a high feature-to-sample ratio, some studies have pointed out that ensemble learning is an effective countermeasure (Awada *et al*., 2012; Saeys *et al*., 2007; He and Yu, 2010). For example, Abeel et al. combined ensemble learning with linear SVM-RFE to successfully improve the robustness and prediction accuracy of selected biomarkers (Abeel *et al*., 2010). Yang and Mao proposed MCF-RFE (multi-criterion fusion-based recursive feature elimination), which outperformed simple SVM-RFE in terms of robustness and prediction accuracy (Yang and Mao, 2011). However, apart from these studies, the application of ensemble learning on gene feature selection is still quite limited, and the effect of different ensemble approaches requires further investigation (Ang *et al*., 2016). We therefore combined ensemble learning with the systems biology feature selector to select genes that have robust prognostic predictive power while also providing biological insights. Furthermore, a comprehensive analysis was carried out to systematically evaluate the results obtained by different ensemble approaches.

Complex diseases such as breast cancer are unlikely caused by the aberration of a single gene but rather by the accumulated distortion of multiple genes, which causes the degradation of a whole biological process that then leads to cancer (Staiger *et al*., 2012). Traditionally, however, the expression of an identified gene biomarker would be directly used to infer the prognosis status. Potential interactions between multiple disease-contributing genes cannot be considered in such a single-gene approach. In contrast, a multi-gene approach would be able to model a complex disease more comprehensively by taking into consideration the expression patterns of multiple genes. Machine learning classifiers can be used for this exact purpose, merging multiple input features into a final prediction. Among various classification models, support vector machine (SVM) and random forest (RF) are common powerful classifiers. Deep neural network (DNN) is also a powerful classification model with high expressivity and has the ability to provide high-level abstract representation of input information. There are successful examples of applying these machine learning models in cancer diagnosis, where gene expression or clinical information is used to predict whether a patient has cancer or not with very high accuracy (Díaz-Uriarte and Alvarez de Andrés, 2006; Akay, 2009). Prognosis prediction, on the other hand, is a much more complicated problem. There are more interacting factors, either known or unknown, which all contribute to the final outcome. We therefore integrated machine learning models with ensemble systems biology feature selection in this study, aiming to predict breast cancer prognosis statuses with multiple genes robustly identified through an ensemble approach.

## 2 Methods

### 2.1 Dataset

The data used in this study is the METABRIC dataset (Curtis *et al*., 2012; Pereira *et al*., 2016) from cBioPortal, which is the largest open-access breast cancer cohort that includes both gene expression data, clinical information, and long-term survival follow-ups. The survival information was used to define the label (prognosis status) for each patient, whereas the gene expression and clinical information were used as model inputs to predict the prognosis status. Although there are other available datasets with gene expression measured by the more popular RNA-Seq technique, either the sample size is too small for relevant analysis or the clinical/survival information is missing.

We defined the label of each patient according to his/her 5-year disease-specific survival (DSS) outcome. For those who died of breast cancer within 5 years, we defined them to be the “poor prognosis class”; For those who died of breast cancer after 5 years, we defined them to be the “good prognosis class”. This binary prognosis status was the label for subsequent classification tasks.

Originally, there were 1980 samples (patients) in the dataset. After excluding those without gene expression data, there were 1904 in total. Among them, 1282 were censored samples, that is, the subject died of another cause or was still alive. Since these cases cannot be labeled as good or poor, we defined these samples as the unlabeled set. For the rest (622) of the labeled samples, we excluded 40 of those without complete clinical information and defined the 582 remaining samples as the labeled set. We then stratified split the labeled set to form a training set (465 samples) and a hold-out testing set (117 samples). Details regarding preprocessing and data distribution can be found in Supplementary A

### 2.2 Systems biology feature selector

The core feature selector used in this study is the systems biology feature selector (Wang *et al*., 2011; Lai *et al*., 2019) (Supplementary B). It is an unsupervised gene feature selector that ranks the importance of genes through interaction network analysis. Based on a prognosis-relevant split criterion, the selector divides samples into two prognosis-distinct groups. Irrelevant genes would be eliminated by ANOVA (analysis of variance) and a gene interaction network is constructed for each group based on BioGrid. The PRV (prognosis relevant value) for each gene is then calculated to summarize how differently a gene interacts with all its partners in two prognosis-distinct interaction networks. Gene feature selection was performed by ranking the genes based on the calculated PRVs (Fig. 1).

**Fig. 1.**
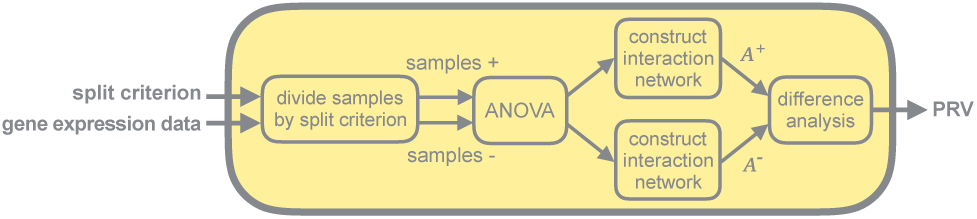
Systems biology feature selector. The required inputs for the systems biology feature selector are a prognostic-relevant split criterion and unlabeled samples with gene expression value. The output is the PRV for each gene feature, which was used to rank and select the genes.

It should be noted that when a different split criterion is assigned to the systems biology feature selector, a different result will be produced and hence can be seen as a different feature selection function. In this study, seven prognosis-relevant split criteria were employed. Five of them were the statuses of well-established breast cancer biomarkers, namely ER, PR, HER2, MKI67, and PLAU, whereas two of them were breast cancer sub-types, specifically the triple negative subtype (TN) and hormone receptor positive subtype (HP).

### 2.3 Ensemble feature selection

In this study, we combined the concept of ensemble learning (Zhang and Ma, 2012) with the systems biology feature selector to improve the robustness of gene feature selection. There are generally two major approaches to ensemble feature selection — data perturbation and function perturbation (He and Yu, 2010). In the data-perturbation ensemble approach, a feature selector is trained multiple times on different sample subsets, resulting in multiple different feature selection outcomes (Fig. 2a).

**Fig. 2.**
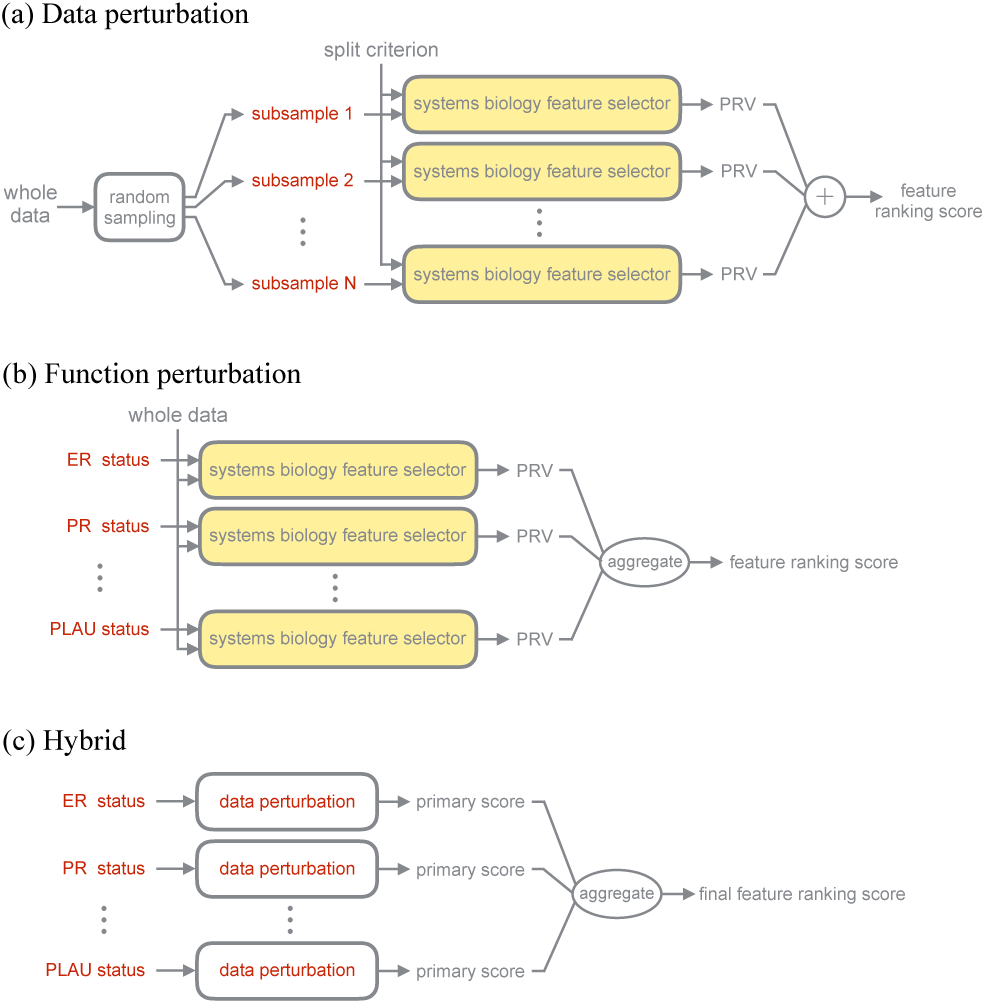
Ensemble feature selection workflow. (a) In data perturbation, multiple sample subsets were generated through random sampling. The systems biology feature selector was trained on different sample subsets. The output scores were then summed together to produce the final score. (b) In function perturbation, different systems biology feature selection functions were all trained on whole training data then aggregated to produce the final score. (c) In the hybrid ensemble approach, different systems biology feature selection functions first underwent data perturbation and then the data perturbation output of different functions were aggregated to produce the final score.

The outcomes are then aggregated together to handle the selection instability with respect to sampling variation. On the other hand, the function-perturbation ensemble approach tries to run different feature selectors on the same dataset then aggregates the outcomes (Fig. 2b). The idea is to capitalize on the strengths of different feature selection algorithms to obtain a robust final output.

Based on the above approaches, a third approach emerged — the hybrid ensemble approach. The hybrid ensemble approach intuitively tries to combine the strengths of data perturbation and function perturbation to further improve the robustness (Fig. 2c). Based on a previous review, detailed research regarding the hybrid ensemble approach is still lacking and there are no previous examples that apply the hybrid ensemble approach on gene feature selection (Awada *et al*., 2012). Therefore, in this study, we comprehensively analyzed the three ensemble approaches mentioned above in the area of gene feature selection.

### 2.4 Prognosis prediction

We trained classifiers to predict the prognosis statuses of patients. In the first stage, the purpose was to evaluate and compare different feature selection methods. Therefore, logistic regression was adopted to directly reflect the performance of selected genes. In the second stage, however, the purpose became finalizing a classifier that produces the best predictive performance. Therefore, more complex classifiers were adopted, including support vector machine (SVM), random forest (RF), and deep neural network (DNN).

A special bimodal structure (Ngiam *et al*., 2011) (Supplementary C) for the DNN was used when combining heterogeneous inputs of gene expression and clinical information. The two data sources were first processed by two separated subnetworks then merged together. This bimodal structure was shown to outperform simple fully connected DNNs (Lai *et al*., 2019).

### 2.5 Evaluation

We used AUC (area under receiver operating characteristic curve) (Hanley and McNeil, 1982) as the main metric for the evaluation of predictive performance, since it provides a comprehensive overview of the performance of the model at all possible classification thresholds.

In the first stage, we performed random validation 100 times to evaluate the stability of a feature selection method. Each time, random validation was carried out by sub-dividing the training set into a smaller training set (3/4) and a validation set (1/4). We evaluated the performance of a feature selection method by focusing on its top-50 ranked genes, and a curve corresponding to the validation AUC of the top-1 ranked gene to the top-50 ranked genes was plotted. We then quantified the overall performance of a feature selection method by the “area” under this top-50 AUC curve. After random validation was performed 100 times, we generated 100 curves and 100 summarized areas. The distribution of these summarized areas was presented with box plot and the averaged curve of 100 curves was also presented to display the rough performance pattern of the top-50 ranked genes. We focused on the top-50 selected genes since, in the case of this study, important genes are usually ranked within the top 50, and a peak performance can be achieved within this window. Genes ranked outside the top 50 add very minor improvement to the predictive performance, and hence it makes less sense to include them when calculating the summarized area. When comparing two feature selection methods, we used the one-tailed paired *t*-test to compare two sets of area distribution. This enabled us to statistically verify if a set of selected genes leads to significantly better predictive performance on different unseen data (100 random validations), which confers a more robust feature selection result.

In the second stage, we used 4-fold cross validation to determine the hyperparameter of our final proposed model. We did not adopt the 100-random validation procedure as in the first stage, since hyperparameter grid search with such random validation setting is not computationally feasible. The averaged performance over 4-fold cross validation was used to present the performance of a hyperparameter set, and the hyperparameter set that led to the highest cross validation performance was selected as the final hyperparameter set. After determining the hyperparameters, we trained the final model with the whole training set with determined hyperparameters and then tested the model on the hold-out test set.

## 3 Results

### 3.1 Comparison of different ensemble approaches

In the first stage, we systematically evaluated the feature selection results of different approaches through 100 random validations as described in Sec. 2.5.

#### 3.1.1 Data-perturbation ensemble approach

The random sampling setting we used in data perturbation was first determined by random validation, in which subsampling 70% of the data each time and repeating five times resulted in the best performance (Supplementary D). We then compared the seven original feature selectors with their data-perturbation versions. The result can be seen in the integrated plot (Fig. 3c-p), and the separated pairwise comparisons for each feature selector are also provided in Supplementary Fig. S3. From Fig. S3, we found that data perturbation improves the robustness in most of the cases except for PR-selector. The improvement was verified through the one-tailed paired *t*-test, which implied that the “summarized area” distribution of the data-perturbation results for ER, HER2, TN, HP, MKI67, and PLAU-selectors were all significantly higher than their corresponding original feature selection results.

**Fig. 3.**
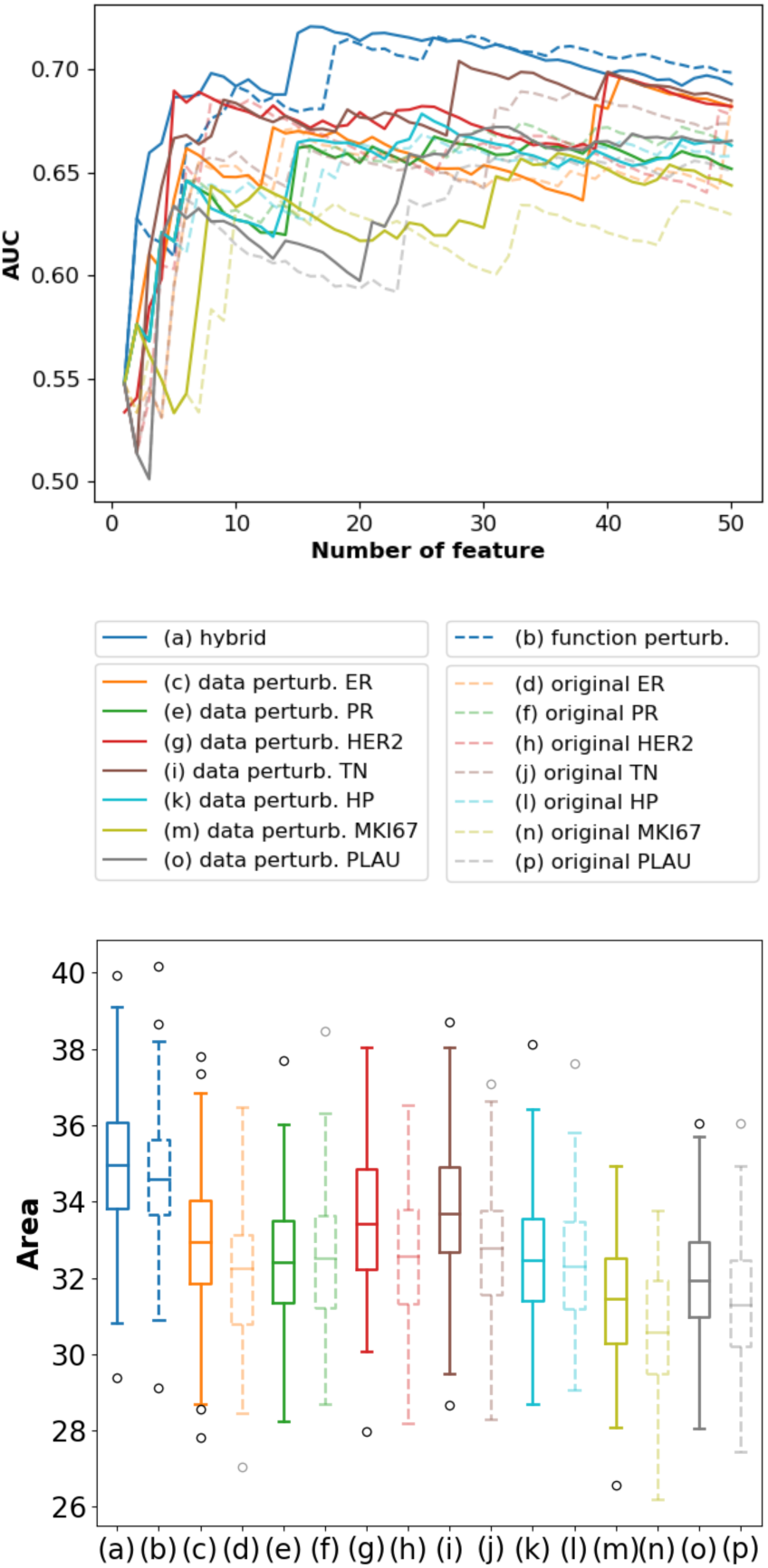
Comparison of different ensemble approaches. (a)-(p) The random validation results of different feature selection approaches are presented. The curves in the left represent the averaged validation AUC for the top-50 selected genes by different approaches. The boxes in the right represent the distribution of the summarized areas under the top-50 curves out of 100 random validations. Higher distribution implies better robustness, since the selected genes have better performance in unseen validation data.

#### 3.1.2 Function-perturbation ensemble approach

Function perturbation aggregates the output score generated by different functions into one final feature ranking score. Other than simply taking the summation, there are many possible aggregation strategies (Pes *et al*., 2017). Through random validation, we found that the rank-mean strategy led to the best performance, by transforming the output scores of seven feature selectors into ranking lists first and then taking the average ranking as the final score (Supplementary E). Having determined the aggregation strategy, we compared the original results of the seven feature selectors (Fig. 3d, f, h, j, l, n, p) with their function-perturbation results (Fig. 3b). A dedicated plot is also provided in Supplementary Fig. S5. Through Fig. S5, we found that function perturbation brought even more significant improvement to the original feature selection results, which was also statistically verified by the one-tailed paired *t*-test.

#### 3.1.3 Hybrid ensemble approach

We further compared the results of function perturbation (Fig. 3b) and data perturbation (Fig. 3c, e, g, i, k, m, o) with the hybrid ensemble approach (Fig. 3a). We found that the hybrid ensemble approach produced the most robust feature selection results among all approaches tested. The improvement was also verified by the one-tailed paired *t*-test. This implies that the genes selected by the hybrid ensemble approach had a consistently better performance in 100 random validations, which is therefore a more robust feature selection result compared to either the result of data perturbation, function perturbation, or the original systems biology feature selector.

As a result, we adopted the best-performing hybrid ensemble approach to select the final gene set. As observed from the top-50 curve of the hybrid ensemble approach (Fig. 3a curve plot), the first 16 genes alone produced the peak performance. Therefore, the first 16 genes were the final gene set we selected, which served as an extension to the inputted prior breast cancer knowledge of well-established biomarkers and subtypes. With much fewer number of features, the 16 final selected genes significantly outperformed the combination of all genes before feature selection (24,338 candidate genes) in random validation (Fig. S6).

### 3.2 Test performance evaluation of final prognosis models

After filtering out genes with the most robust prognosis predictive power, we moved on to finalizing the prognosis classification model in the second stage. Rather than the simple logistic regression as used in the first stage, more complex models such as SVM, RF, and DNN were considered to construct the final prognosis models. The hyperparameters were determined through 4-fold cross validation, listed in Supplementary G. After determining the hyperparameters, the final models were trained with whole training data and tested on the hold-out test set. Considering that both gene expression data and clinical information might not always be available at the same time, we proposed different models with only gene expression input, only clinical information input, and combined input.

Firstly, models with only gene features achieved an AUC between 0.7443 and 0.7672 (Table 1a). The input features are the corresponding genes of well-established breast cancer biomarkers (ESR1, PGR, ERBB2, MKI67, PLAU) and the final selected genes in Sec. 3.1.3. Through the test performance, we found that the expression pattern of these selected genes alone can give an accurate prediction towards the prognosis status.

**Table 1.** Test performance evaluation of final models

|  | (a) Gene |  | (b) Clinical |  | (c) Combined |  |
| --- | --- | --- | --- | --- | --- | --- |
|  | Accuracy | AUC | Accuracy | AUC | Accuracy | AUC |
| SVM | 0.6838 | 0.7443 | 0.6154 | 0.6657 | 0.6752 | 0.7677 |
| RF | 0.6923 | 0.7663 | **0.6581** | **0.6850** | **0.7265** | 0.7815 |
| DNN | **0.7009** | **0.7672** | 0.6154 | 0.6833 | 0.7179 | **0.7836** |

Secondly, models with only clinical features achieved an AUC between 0.6657 and 0.6850 (Table 1b). The input features are the 10 clinical features listed in Supplementary A. Through pairwise comparison of the first two columns in Table 1, we found that gene feature models performed substantially better than clinical feature models. This implies that the selected genes can reflect the prognosis status more directly than typical clinical features which are usually thought to be the most directly linked to the prognosis status. However, under the circumstances in which gene expression measurements are not available, the predicted prognosis by clinical feature models can still serve as a reference.

Finally, the models combining both gene and clinical features achieved an AUC between 0.7677 and 0.7836 (Table 1c). The structure for the DNN we used here is the bimodal structure as described in Supplementary C. We found that bimodal DNN successfully combined heterogeneous inputs of gene expression and clinical information, achieving the highest AUC among all models.

We further validated the performance of bimodal DNN through traditional survival analysis. The concordance index (CI) (Harrell, 2015) of the bimodal DNN was 0.6683, which outperformed the traditional cox model (Cox, 1972; Bradburn *et al*., 2003) trained with the same input features (CI = 0.6251). In addition, the survival curve (Clark *et al*., 2003; Harrell, 2015) of the good and poor prognosis groups predicted by bimodal DNN is illustrated in Fig. 4. As observed from the plot, after five years, the overall survival rate of the predicted good prognosis group is 0.68, while that of the predicted poor prognosis group is only 0.24. A log-rank test (Peto *et al*., 1977; Clark *et al*., 2003) also showed that the survival rate of two groups of patients is significantly different (*p*-value = 1.763×10^−5^).

**Fig. 4.**
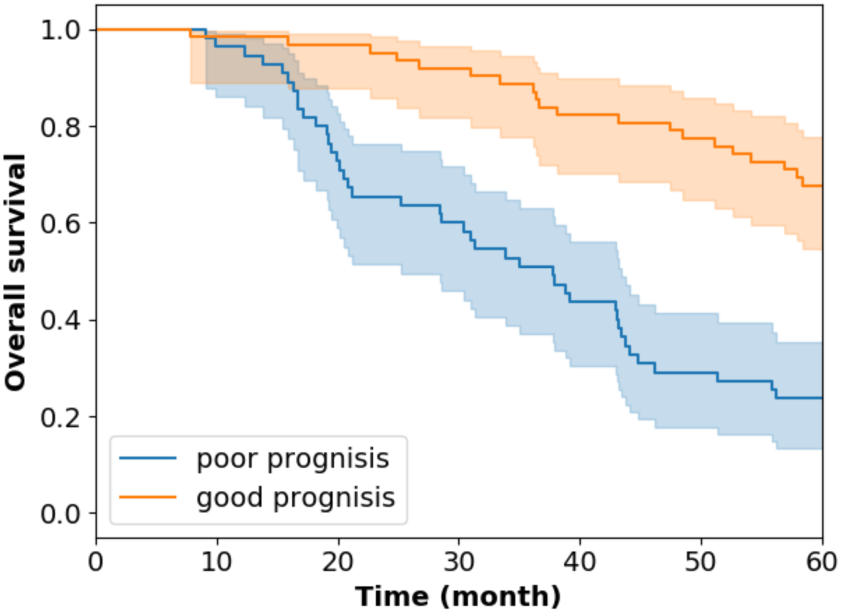
Kaplan-Meier plot of two groups of patients classified by bimodal DNN. The blue curve represents the overall survival rate over time for the poor prognosis group of patients predicted by bimodal DNN. The orange curve represents the good prognosis group predicted by bimodal DNN.

## 4 Discussion

Selecting robust gene features has long been a challenging issue due to the high dimensionality and low sample size properties of microarray data. To address the problem, we introduced ensemble learning into our systems biology feature selection pipeline. We systematically evaluated three ensemble approaches through 100 random validations, which is one of the first comprehensive analyses of different ensemble approaches on gene feature selection. The results show that all three ensemble approaches improved the feature selection robustness. Among all, the hybrid ensemble approach resulted in the most significant improvement, such that the selected genes achieved the highest overall performance on different validation sets. In addition, while the most popular data-perturbation ensemble approach does bring improvement, the less frequently used function-perturbation ensemble approach can actually bring about more significant improvement with just a few numbers of ensembles.

Further analysis on function perturbation showed that the final aggregation can benefit even from adding suboptimal feature selection functions. Initially, only ER, PR, and HER2 were adopted as split criteria, since they are the most high-confidence, well-established breast cancer biomarkers. TN and HP are major prognosis-relevant subtypes, but individually, they did not outperform the primary function perturbation (ER + PR + HER2; Fig. 5a). However, adding the suboptimal feature selectors TN and HP to the primary function perturbation improved the performance surprisingly (Fig. 5e). Similarly, when we further aggregated MKI67 and PLAU, the performance boosted again (Fig. 5h), which then became the final version of function perturbation. This indicates that by merging a few suboptimal-but diverse-functions, function perturbation can achieve significantly better performance.

**Fig. 5.**
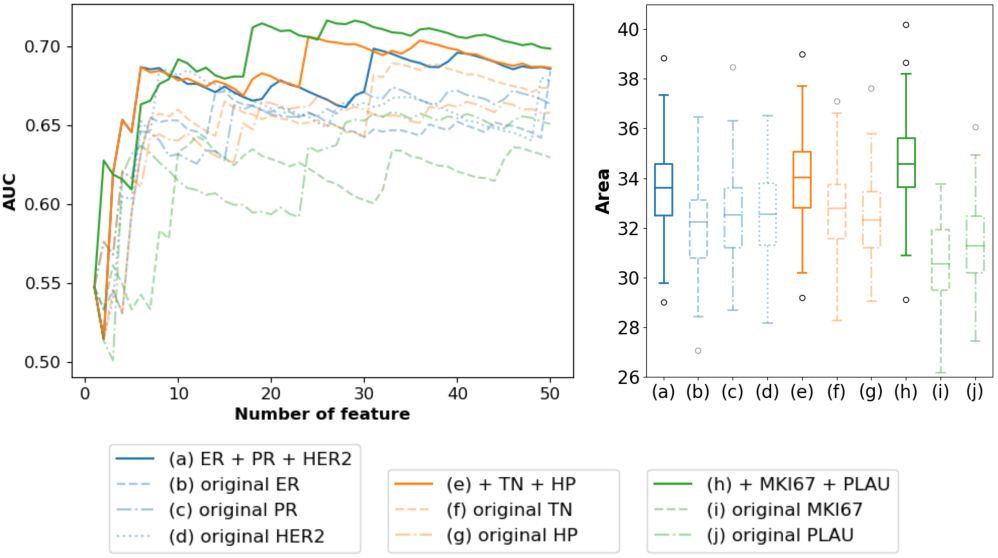
Aggregating a different number of functions in function perturbation. (a) Aggregating ER, PR, and HER2-selector. (b)-(d) Original feature selection result of ER, PR, and HER2-selector. (e) Further aggregating TN and HP. (f)-(g) Original feature selection result of TN and HP-selector. (h) Further aggregating MKI67 and PLAU. (i)-(j) Original feature selection result of MKI67 and PLAU-selector.

On the other hand, although compared to function perturbation, data perturbation brings relatively minor robustness improvement, both approach further improve upon each other. The highest performance was achieved only in the final aggregation of data diversity and function diversity in the hybrid ensemble approach. Therefore, the conclusion that we draw from random validation analysis is that, when computational resource is limited, function perturbation would be recommended over data perturbation. However, when computational resources are not the major concern, hybrid ensemble approach would be the best strategy to ensure robustness.

Due to the core systems biology feature selector that was wrapped in the ensemble learning workflow, our feature selection method also successfully incorporated prior biological knowledge to select genes that provide biological insights. Firstly, STRING interaction network analysis (Szklarczyk *et al*., 2017) showed that the 16 selected genes are tightly linked through experimental or literary verified interactions (Fig. 6). Among these genes, ESR1 (Kim *et al*., 2011), ELAVL1 (Yuan *et al*., 2010; López de Silanes *et al*., 2005), EGFR (Masuda *et al*., 2012), and YQHAQ (Santarius *et al*., 2010) were already known to be related to breast cancer. Strong linkage between these well-studied breast-cancer-related genes and other identified genes makes the identified, but under-studied, genes more reasonable targets for further experimental investigation.

**Fig. 6.**
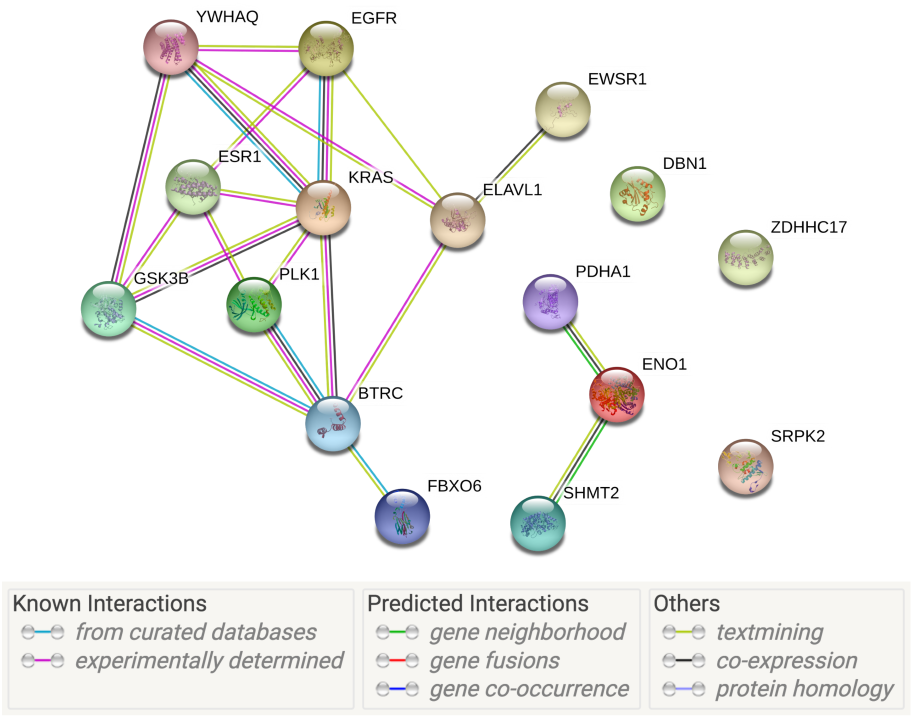
STRING analysis of 16 selected genes.

Secondly, the systems biology feature selector identifies important genes based on interaction network analysis. Therefore, compared to pure statistical approaches that typically focus on the differential expression of each individual gene between two patient groups, our approach focuses on the topological aspect differences. We conducted enrichment analysis on an expanded list of the top-50 ranked genes by hybrid ensemble approach (listed in Supplementary F) to find out what pathways the identified genes generally fall into. In the resulting list of biological process enrichment analysis, we found that the genes are highly involved in pathways such as cell cycle and ubiquitination. For example, highly ranked genes such as BTRC (#2), FBXO6 (#3), SHMT2 (#4), GSK3B (#16), FBXW7 (#18), and UCHL5 (#19) are related to ubiquitination. It is known that the misregulated expression of E3 ubiquitin ligases contributes to aberrant oncogenic signaling (Gallo *et al*., 2017), where FBXW7 is an example. FBXW7 is a component of the SCF (SKP1, CUL-1, F-box protein) E3 ubiquitin ligase complex, where its down-regulation in breast, colorectal, gastric, and cholangiocarcinoma (CCA) tumors correlates with poor prognosis and survival, elevated tumor invasion, and occurrence of metastasis (Iwatsuki *et al*., 2010; Yang *et al*., 2015; Ibusuki *et al*., 2011). We therefore think that other identified ubiquitination-related genes in the list may link to similar breast cancer pathogenesis mechanism as well. In addition, it is known that ubiquitination pathways are potential druggable pathways (Gallo *et al*., 2017). Thus, the genes selected in this study may also be potential druggable targets. On the other hand, among the top-50 list, there are also genes related to cell cycle, such as PLK1 (#14), AURKA (#21), CDK4 (#31), and CDK1 (#43). For example, CDK1 and CDK4 play key roles in cell cycle regulation (Malumbres and Barbacid, 2005). Their over-expression is closely related to proliferative diseases such as cancer (Kim *et al*., 2007). Therefore, CDK1 and CDK4 were found to be potential cancer therapeutic targets. CDK1 is also involved in the activation of AURKA and PLK1 in the complex cell cycle regulatory network, and together they control whether a cell enters the mitosis phase (Asteriti *et al*., 2015; Lindqvist *et al*., 2009). Due to the deterministic role in cell cycle regulation, AURKA and PLK1 are also possible targets for inhibiting abnormal proliferation (Giet *et al*., 2005; Spankuch-Schmitt *et al*., 2002). Disorders in cell-cycle-related pathways have key influences on the prognosis of breast cancer, and our feature selection result highlights that CDK1, CDK4, AURKA, and PLK1 may play particularly important roles in the complex cell cycle regulatory network, which in turns affects breast cancer prognosis.

With the selected gene features that provide biological insights and robust predictive performance, we moved on to finalizing prognosis prediction models in the second stage of the study. Through test performance evaluation, we found that models with gene feature alone can achieve an AUC between 0.7443 and 0.7672. This performance achieved by a multigene approach is higher than the AUC of any component gene as a single biomarker (Supplementary H). This indicates that a multi-gene approach can indeed model the complex molecular process of breast cancer more comprehensively through joint evaluation of multiple genes. On the other hand, clinical feature models can also serve as a reference when gene expression data is not available. However, models that combine both gene expression and clinical information were the ones that achieved the best predictive performance, with bimodal DNN achieving the highest AUC among all. Additional survival analysis also indicates that bimodal DNN can successfully differentiate patients with different prognosis status.

In conclusion, our study demonstrated that ensemble learning can help improve gene feature selection robustness. The selected genes provide insight into the complex breast cancer molecular process from a topological aspect and serve as good targets for further experimental validation. Furthermore, test evaluation and survival analysis showed that bimodal DNN can accurately predict breast cancer prognosis, which would in turn help guide personalized and precise treatment.

## Acknowledgements

We would like to thank Uni-edit (www.uni-edit.net) for editing and proofreading this manuscript.

## Funding

This work has been supported by the Ministry of Technology and Science, Taiwan, under Grants 108-2218-E-007-052.

## Conflict of Interest

none declared.

## References

Abeel, T. et al. (2010) Robust biomarker identification for cancer diagnosis with ensemble feature selection methods. Bioinformatics, 26, 392–398.

Akay, M.F. (2009) Support vector machines combined with feature selection for breast cancer diagnosis. Expert Syst. Appl., 36, 3240–3247.

Ang, J.C. et al. (2016) Supervised, unsupervised, and semi-supervised feature selection: A review on gene selection. IEEE/ACM Trans. Comput. Biol. Bioinforma., 13, 971–989.

Asteriti, I.A. et al. (2015) Cross-Talk between AURKA and Plk1 in Mitotic Entry and Spindle Assembly. Front. Oncol., 5, 283.

Awada, W. et al. (2012) A review of the stability of feature selection techniques for bioinformatics data. Proc. 2012 IEEE 13th Int. Conf. Inf. Reuse Integr. IRI 2012, 356–363.

Bradburn, M.J. et al. (2003) Survival Analysis Part II: Multivariate data analysis-An introduction to concepts and methods. Br. J. Cancer, 89, 431–436.

Carey, L.A. et al. (2007) The Triple Negative Paradox: Primary Tumor Chemosensitivity of Breast Cancer Subtypes. Clin. Cancer Res., 13, 2329–2334.

Clark, T.G. et al. (2003) Survival Analysis Part I: Basic concepts and first analyses. Br. J. Cancer, 89, 232–238.

Cox, D.R. (1972) Regression Models and Life-Tables. J. R. Stat. Soc. Ser. B, 34, 187–220.

Curtis, C. et al. (2012) The genomic and transcriptomic architecture of 2,000 breast tumours reveals novel subgroups. Nature, 486, 346–352.

Dent, R. et al. (2007) Triple-Negative Breast Cancer: Clinical Features and Patterns of Recurrence. Clin. Cancer Res., 13, 4429–4434.

Díaz-Uriarte, R. and Alvarez de Andrés, S. (2006) Gene selection and classification of microarray data using random forest. BMC Bioinformatics, 7, 3.

Duffy, M.J. et al. (2017) Clinical use of biomarkers in breast cancer: Updated guidelines from the European Group on Tumor Markers (EGTM). Eur. J. Cancer, 75, 284–298.

Dunnwald, L.K. et al. (2007) Hormone receptor status, tumor characteristics, and prognosis: a prospective cohort of breast cancer patients. Breast Cancer Res., 9, R6.

Gallo, L.H. et al. (2017) The importance of regulatory ubiquitination in cancer and metastasis. 16, 634–648.

Giet, R. et al. (2005) Aurora kinases, aneuploidy and cancer, a coincidence or a real link? Trends Cell Biol., 15, 241–250.

Hanley, J.A. and McNeil, B.J. (1982) The meaning and use of the area under a receiver operating characteristic (ROC) curve. Radiology, 143, 29–36.

Harrell, F.E. (2015) Regression Modeling Strategies.

He, Z. and Yu, W. (2010) Stable feature selection for biomarker discovery. Comput. Biol. Chem., 34, 215–225.

Ibusuki, M. et al. (2011) Reduced expression of ubiquitin ligase FBXW7 mRNA is associated with poor prognosis in breast cancer patients. Cancer Sci., 102, 439–445.

Iwatsuki, M. et al. (2010) Loss of FBXW7, a cell cycle regulating gene, in colorectal cancer: Clinical significance. Int. J. Cancer, 126, 1828–1837.

Kalousis, A. et al. (2007) Stability of feature selection algorithms: a study on high-dimensional spaces. Knowl. Inf. Syst., 12, 95–116.

Kim, C. et al. (2011) Estrogen receptor (ESR1) mRNA expression and benefit from tamoxifen in the treatment and prevention of estrogen receptor-positive breast cancer. J. Clin. Oncol., 29, 4160–7.

Kim, S.-Y. (2009) Effects of sample size on robustness and prediction accuracy of a prognostic gene signature. BMC Bioinformatics, 10, 147.

Kim, S.J. et al. (2007) Determination of the specific activity of CDK1 and CDK2 as a novel prognostic indicator for early breast cancer. Ann. Oncol., 19, 68–72.

Lai, Y.-H. et al. (2019) Predicting the prognosis of non-small cell lung cancer by integrating microarray and clinical data with deep learning. bioRxiv, 656140.

Lehmann, B.D. et al. (2011) Identification of human triple-negative breast cancer subtypes and preclinical models for selection of targeted therapies. J. Clin. Invest., 121, 2750–2767.

Lindqvist, A. et al. (2009) The decision to enter mitosis: feedback and redundancy in the mitotic entry network. J. Cell Biol., 185, 193–202.

López de Silanes, I. et al. (2005) HuR: post-transcriptional paths to malignancy. RNA Biol., 2, 11–3.

Malumbres, M. and Barbacid, M. (2005) Mammalian cyclin-dependent kinases. Trends Biochem. Sci., 30, 630–641.

Masuda, H. et al. (2012) Role of epidermal growth factor receptor in breast cancer. Breast Cancer Res. Treat., 136, 331–345.

Ngiam, J. et al. (2011) Multimodal deep learning. In, Proceedings of the 28th international conference on machine learning (ICML-11)., pp. 689–696.

Pereira, B. et al. (2016) The somatic mutation profiles of 2,433 breast cancers refines their genomic and transcriptomic landscapes. Nat. Commun., 7.

Pes, B. et al. (2017) Exploiting the ensemble paradigm for stable feature selection: A case study on high-dimensional genomic data. Inf. Fusion, 35, 132–147.

Peto, R. et al. (1977) Design and analysis of randomized clinical trials requiring prolonged observation of each patient. II. Analysis and examples. Br. J. Cancer, 35, 1–39.

Polyak, K. (2011) Heterogeneity in breast cancer. J. Clin. Invest., 121, 3786–8.

Saeys, Y. et al. (2007) A review of feature selection techniques in bioinformatics. Bioinformatics, 23, 2507–2517.

Santarius, T. et al. (2010) A census of amplified and overexpressed human cancer genes. Nat. Rev. Cancer, 10, 59–64.

Spankuch-Schmitt, B. et al. (2002) Effect of RNA Silencing of Polo-Like Kinase-1 (PLK1) on Apoptosis and Spindle Formation in Human Cancer Cells. JNCI J. Natl. Cancer Inst., 94, 1863–1877.

Staiger, C. et al. (2012) A critical evaluation of network and pathway-based classifiers for outcome prediction in breast cancer. PLoS One, 7.

Stark, C. et al. (2006) BioGRID: a general repository for interaction datasets. Nucleic Acids Res., 34, D535–D539.

Szklarczyk, D. et al. (2017) The STRING database in 2017: quality-controlled protein–protein association networks, made broadly accessible. Nucleic Acids Res., 45, D362–D368.

Wang, Y.-C. et al. (2011) A network-based biomarker approach for molecular investigation and diagnosis of lung cancer. BMC Med. Genomics, 4, 2.

Yang, F. and Mao, K.Z. (2011) Robust feature selection for microarray data based on multicriterion fusion. IEEE/ACM Trans. Comput. Biol. Bioinform., 8, 1080–1092.

Yang, H. et al. (2015) FBXW7 suppresses epithelial-mesenchymal transition, stemness and metastatic potential of cholangiocarcinoma cells. Oncotarget, 6, 6310–6325.

Yuan, Z. et al. (2010) HuR, a key post-transcriptional regulator, and its implication in progression of breast cancer. Histol. Histopathol., 25, 1331–40.

Zhang, C. and Ma, Y. (2012) Ensemble Machine Learning Zhang, C. and Ma, Y. (eds) Springer US, Boston, MA.

