## Supplementary Information for "Integrating ensemble systems biology feature selection and bimodal deep neural network for breast cancer prognosis prediction"

#### A. Data preprocessing and data distribution

There are 24,338 available gene features and 10 relevant clinical features in the METABRIC dataset.

The gene features are preprocessed microarray gene expression values. Although gene expression measured through the RNA-Seq technique is the major trend nowadays, which contains lower background noise, METABRIC is the largest available open-access breast cancer cohort containing gene expression, clinical information, and long-term survival data that allows relevant data analysis. We did not use a combined dataset by merging multiple small cohorts, since different platforms, sample curation workflows, and experimental procedures can all lead to different measurement outputs. Even if multiple cohorts are combined after careful preprocessing and standardization, it is still difficult to obtain robust feature selection and prognosis prediction results using such a combined dataset with heterogeneous composition.

The clinical features were derived from the raw clinical information in the dataset, which include age, tumor size, neoplasm histologic grade, cellularity, menopausal state, radio therapy, chemotherapy, hormone therapy, breast conserving surgery, and breast mastectomy surgery. Since most features (the latter six) are binary, we normalized the other features (the first four) according to the training data, ensuring that the scale of clinical features did not differ significantly from each other.

After excluding samples with missing values, the dataset was divided into an unlabeled, training, and hold-out testing sets. Table S1 provides a summary of the three subsets.

| | **Unlabeled** | **Training** | **Testing** |
|---|---|---|---|
| **Samples** | 1282 | 465 | 117 |
| **Class** | | | |
| Good prognosis | - | 221 (48%) | 55 (47%) |
| Poor prognosis | - | 244 (52%) | 62 (53%) |
| **Median DSS time (months)** | - | 56.33 | 56.27 |
| **Median age (years)** | 61.99 | 61.55 | 59.82 |
| **Median tumor size (mm)** | 21 | 26 | 25 |
| **ER status** | | | |
| Positive | 1023 (80%) | 326 (70%) | 81 (69%) |
| Negative | 259 (20%) | 139 (30%) | 36 (31%) |
| **PR status** | | | |
| Positive | 729 (57%) | 207 (45%) | 54 (46%) |
| Negative | 553 (43%) | 258 (55%) | 63 (54%) |

| | | | |
|---|---|---|---|
| **HER2 status** | | | |
| Positive | 120 (9%) | 86 (18%) | 23 (20%) |
| Negative | 1162 (91%) | 379 (82%) | 94 (80%) |
| **Menopausal state** | | | |
| Pre | 260 (20%) | 105 (23%) | 32 (27%) |
| Post | 1022 (80%) | 360 (77%) | 85 (73%) |
| **Neoplasm Histologic Grade** | | | |
| 1 | 140 (11%) | 16 (3%) | 7 (6%) |
| 2 | 530 (43%) | 164 (35%) | 38 (32%) |
| 3 | 556 (45%) | 285 (61%) | 72 (62%) |
| **Cellularity** | | | |
| Low | 142 (11%) | 40 (9%) | 15 (13%) |
| Moderate | 474 (38%) | 173 (37%) | 50 (43%) |
| High | 626 (50%) | 252 (54%) | 52 (44%) |

**Table S1. Data distribution overview**

#### B. Systems biology feature selector

The inputted samples were divided into two groups according to the binary split criterion that was assigned, for example, ER+ samples and ER- samples. Genes without significant differential expression between the two groups were excluded by ANOVA. Next, we constructed interaction networks for each group based on the interaction information documented in the BioGRID database. We used gene expression data to estimate the interaction ability between genes and excluded false positive links. The constructed interaction networks would therefore be disease-specific and tailored for the inputted group of samples.

The main assumption of the network construction method is that the expression level of a gene is affected by other genes, which can thus be represented by the linear combination of the expression level of its interaction partners:

$$x_i[n] = \sum_{j \in \mathrm{G}_i} a_{ij} x_j[n] + \varepsilon_i[n] \tag{1}$$

where $x_i[n]$ is the expression level of gene $i$ for patient $n$; $a_{ij}$ is the interaction ability between genes $i$ and $j$; $\mathrm{G}_i$ is the set of genes that are related to gene $i$ according to BioGRID; and $\varepsilon_i[n]$ is stochastic noise. Equation (1) can be rewritten into the matrix form:

$$\boldsymbol{X} = \boldsymbol{AX} + \boldsymbol{E} \tag{2}$$

where

$$\boldsymbol{X} = \begin{bmatrix} x_1[1] & \cdots & x_1[N] \\ \vdots & \ddots & \vdots \\ x_M[1] & \cdots & x_M[N] \end{bmatrix}, \boldsymbol{A} = \begin{bmatrix} a_{11} & \cdots & a_{1M} \\ \vdots & a_{ij} & \vdots \\ a_{M1} & \cdots & a_{MM} \end{bmatrix}, \boldsymbol{E} = \begin{bmatrix} \varepsilon_1[1] & \cdots & \varepsilon_1[N] \\ \vdots & \ddots & \vdots \\ \varepsilon_M[1] & \cdots & \varepsilon_M[N] \end{bmatrix}$$

where $M$ is the number of genes left after excluding those without differential expression by ANOVA, and $N$ is the sample size. The interaction abilities were estimated by LMMSE (Linear Minimum Mean Square Error). Afterwards, we performed model selection and excluded false positive interactions through AIC (Akaike information criterion) and *t*-test. If $a_{ij}$ is not equal to $a_{ji}$, we took the one with the larger absolute value as the final interaction ability between genes $i$ and $j$. After calculating all the interaction abilities, we then obtain the final interaction ability matrix A. By constructing interaction networks for two sample groups (e.g., ER+ samples and ER- samples), we would get $A^+$ and $A^-$ for each group. We define the difference matrix D to be:

$$\boldsymbol{D} = \boldsymbol{A}^{+} - \boldsymbol{A}^{-} = \begin{bmatrix} d_{11} & \cdots & d_{1M} \\ \vdots & \ddots & \vdots \\ d_{M1} & \cdots & d_{MM} \end{bmatrix} = \begin{bmatrix} a_{11}^{+} - a_{11}^{-} & \cdots & a_{1M}^{+} - a_{1M}^{-} \\ \vdots & \ddots & \vdots \\ a_{M1}^{+} - a_{M1}^{-} & \cdots & a_{MM}^{+} - a_{MM}^{-} \end{bmatrix} \tag{3}$$

where $d_{ij}$ is the difference in interaction ability between genes $i$ and $j$. The prognosis relevance value (PRV) is then defined as:

$$PRV_i = \sum_{j=1}^{M} |d_{ij}| \tag{4}$$

which is the summarized interaction ability difference between gene $i$ and its interaction partners. For each gene, a higher PRV implies greater difference in its interaction abilities between two networks. Since the two networks represent different prognosis statuses, genes with high PRVs can be selected as potential prognosis biomarkers, which serve as an extension to the original inputted prognosis-relevant split criterion.

### C. Bimodal DNN

Figure S1 illustrates the structure of bimodal DNN. The bimodal DNN processes gene expression input and clinical input with two separated subnetworks. The output of two subnetworks were then merged together, processed by successive hidden layers, and then turned into a final prediction output. The weights of the subnetworks were pre-trained. During the first phase of training, we froze the weights of subnetworks and trained only the weights between the merged layer and final output. During the second phase of training, all weights were unfrozen to allow fine-tuning of the whole network.

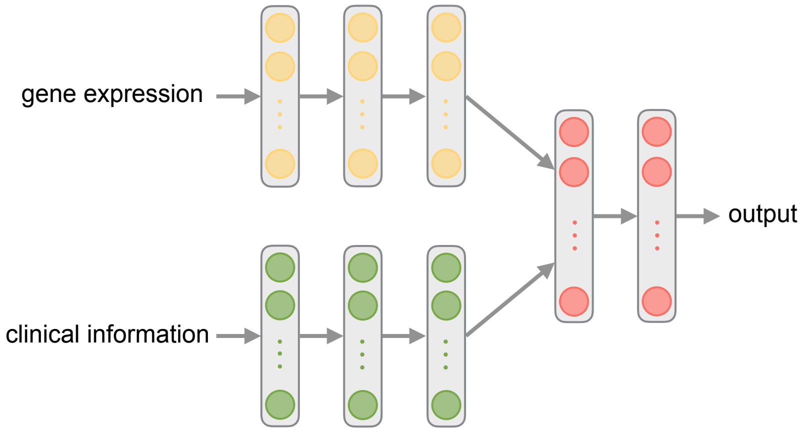


**Fig. S1. Bimodal DNN model structure**

#### D. Data perturbation ensemble approach

For the data perturbation random sampling setting, we tried subsampling 90%, 80%, and 70% of the whole data each time. In addition, we tried different number of subsamples: repeating 5, 10, 20, or 30 times. There were therefore 12 possible combinations of subsampling rates and number of subsamples. We evaluated the 12 settings through random validation by adding the areas of seven feature selectors to be the final summarized area.

From the comparison (Fig. S2), we found that no matter what setting, all of the data perturbation results outperformed those of the original feature selection. Among all, subsampling 70% and repeating 5 times achieved the highest performance, which became the final data perturbation setting we used.

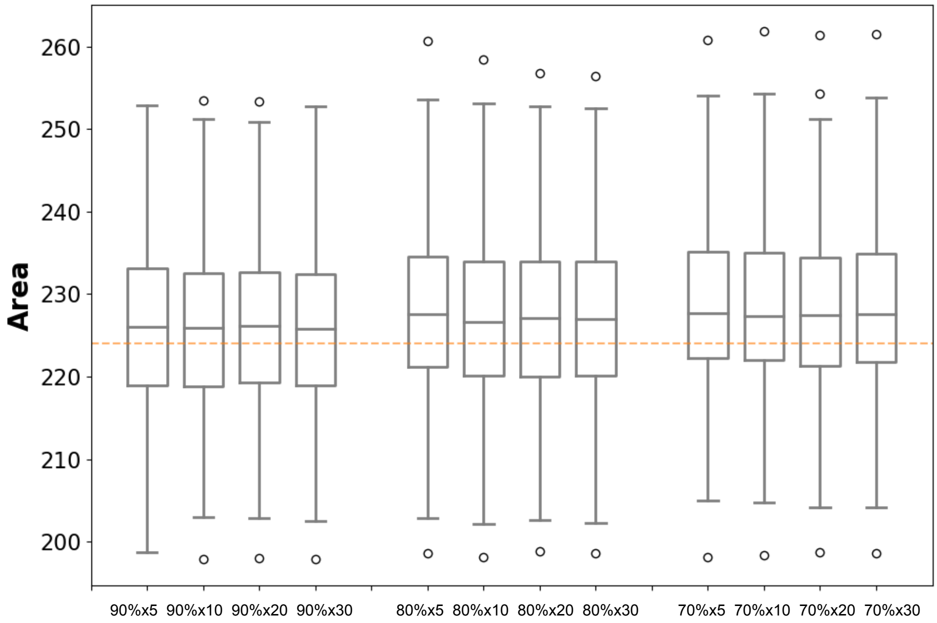


**Fig. S2. Comparison of different data perturbation subsampling settings.** The orange line represents the median of the original feature selection result.

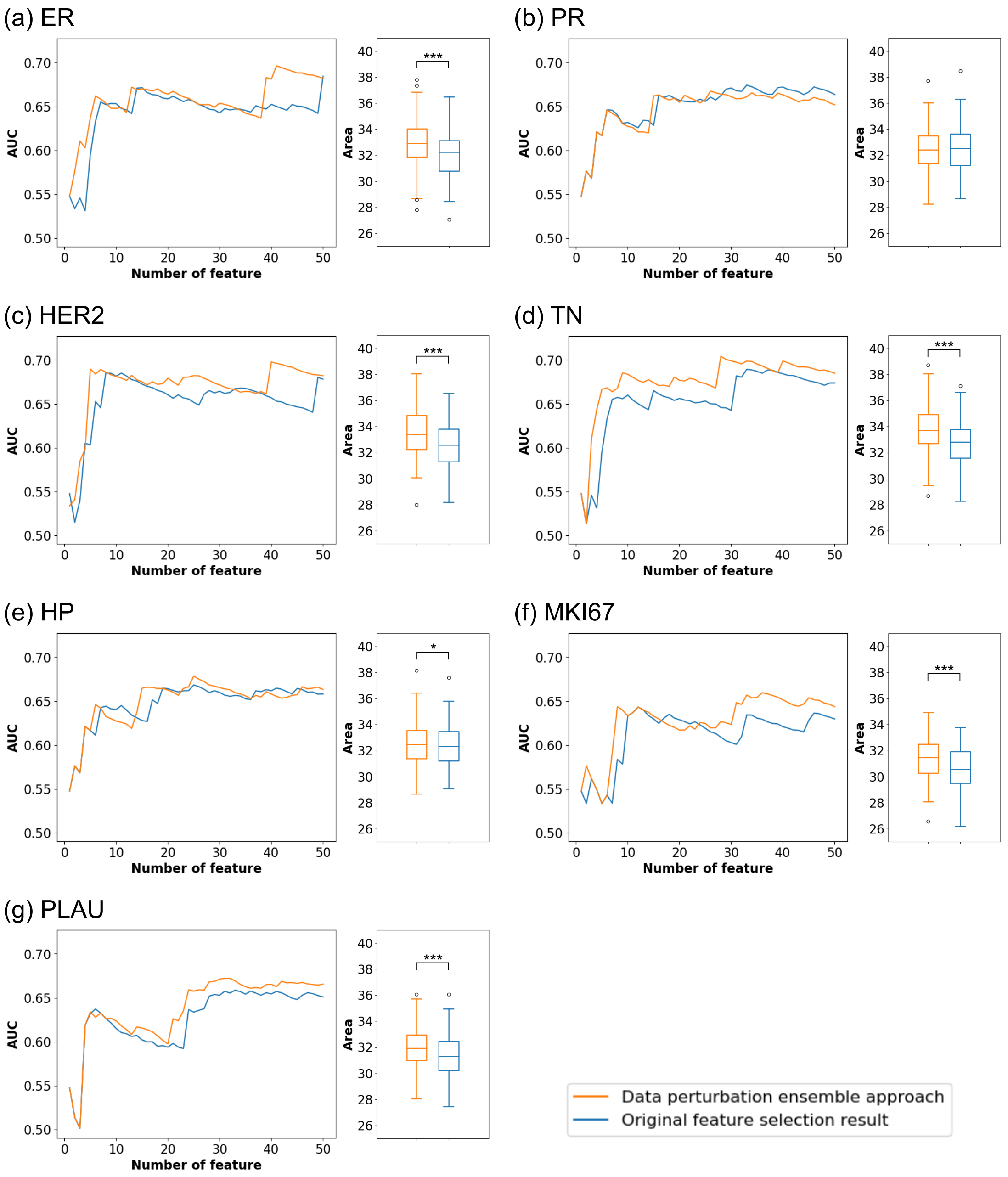


**Fig. S3. Pairwise comparison of data perturbation and original feature selection results**

#### E. Function perturbation ensemble approach

Since the output scale of different systems biology feature selection functions are different, we carefully evaluated the aggregation strategy when combining multiple PRV lists in function perturbation. The following are the aggregation strategies we tested:

(1) **PRV-median**: For each gene, take the median in all PRV lists as its final feature ranking score.
(2) **PRV-norm-median**: Normalize each component PRV list to 0–1. For each gene, take the median of all normalized PRV lists as its final feature ranking score.
(3) **PRV-mean**: For each gene, take the mean in all PRV lists as its final feature ranking score.
(4) **PRV-norm-mean**: Normalize each component PRV list to 0–1. For each gene, take the mean in all normalized PRV lists as its final feature ranking score.
(5) **Rank-median**: Transform each component PRV list into ranking list. For each gene, take the median ranking in all lists as its final feature ranking score.
(6) **Rank-mean**: Transform each component PRV list into ranking list. For each gene, take the mean ranking of all lists as its final feature ranking score.
(7) **Top N overlap**: Take the interaction of the top N genes from each list as the final selected gene. This aggregation strategy cannot output a final feature ranking score but just final selected genes.

Through random validation (Fig. S4) we found that rank-mean and top N overlap produce the best performance. However, since the top N overlap strategy cannot output a feature ranking score and the number of final selected genes cannot be determined by the user, there will be more limitation when using the top N overlap strategy in real applications. Therefore, we adopted rank-mean as our final aggregation strategy for function perturbation.

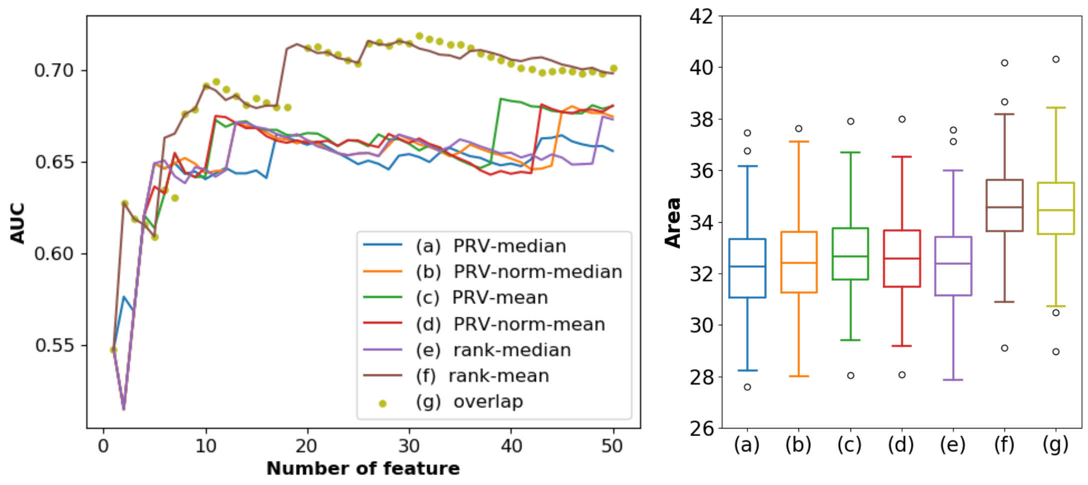


**Fig. S4. Comparison of different function perturbation aggregation strategies**

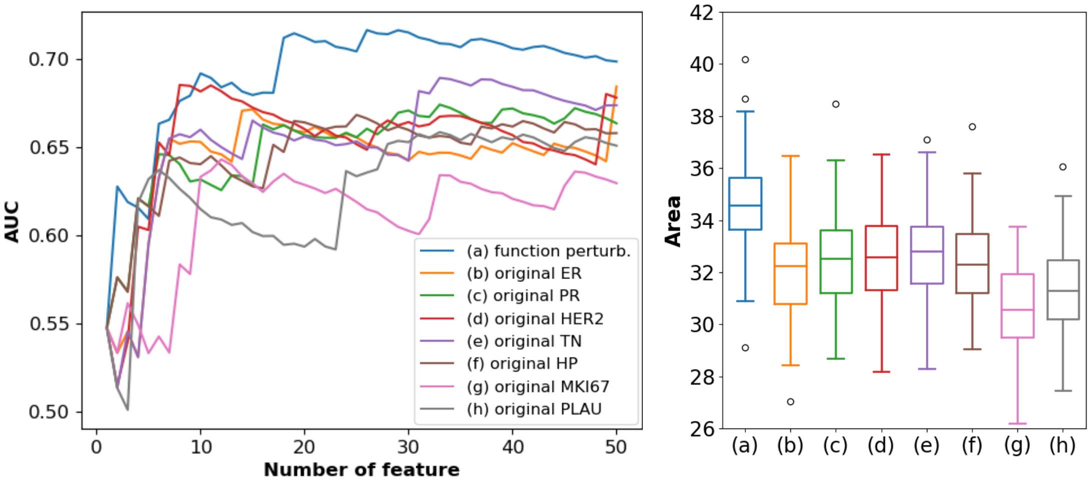


**Fig. S5. Comparison of function perturbation and original feature selection results**

### F. The final selected gene set

The top 50 selected genes via the hybrid ensemble approach are listed below. The final selected genes are the first 16 genes that produced the peak performance.

| | | | | | | | | | |
|---|---|---|---|---|---|---|---|---|---|
| 1 | ELAVL1 | 11 | ZDHHC17 | 21 | AURKA | 31 | CDK4 | 41 | EIF4E2 |
| 2 | EGFR | 12 | ENO1 | 22 | ARRB1 | 32 | SLC15A1 | 42 | IGBP1 |
| 3 | BTRC | 13 | DBN1 | 23 | FLNA | 33 | MCM5 | 43 | CDK1 |
| 4 | FBXO6 | 14 | PLK1 | 24 | CREBBP | 34 | PPP2CA | 44 | CHEK1 |
| 5 | SHMT2 | 15 | ESR1 | 25 | PCM1 | 35 | TCTN1 | 45 | CTDSPL |
| 6 | KRAS | 16 | GSK3B | 26 | ANLN | 36 | TENC1 | 46 | ECT2 |
| 7 | SRPK2 | 17 | HIST1H3A | 27 | RPS14 | 37 | KCTD3 | 47 | AHSA1 |
| 8 | YWHAQ | 18 | FBXW7 | 28 | TRIM23 | 38 | CENPJ | 48 | ACTR2 |
| 9 | PDHA1 | 19 | UCHL5 | 29 | RPL6 | 39 | HDAC11 | 49 | POC5 |
| 10 | EWSR1 | 20 | SYNCRIP | 30 | TUBA1C | 40 | FUS | 50 | YBX1 |

**Table S2. Top 50 selected genes via hybrid ensemble approach**

We compared the performance of the final feature selection result (16 genes) with all genes before feature selection (24,338 genes) through random validation. We found that the selected genes significantly outperform all genes with a much smaller number of features, verified by the one-tailed paired *t*-test. A lower number of features reduces cost in clinical application and prevents model overfitting.

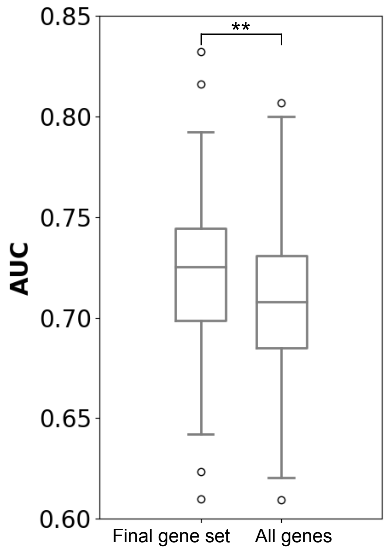


**Fig. S6. Comparison of final selected genes and all genes**

#### G. Hyperparameter grid search result

Below lists the hyperparameter grid search range for SVM, RF, and DNN. We searched through all possible combinations of the listed values. The final determined hyperparameter values are the combinations that achieved the best 4-fold cross validation performance, which are highlighted in bold.

| | Gene feature SVM | Clinical feature SVM | Combined feature SVM |
|---|---|---|---|
| **C** | 0.1, **1**, 10, 100 | 0.1, 1, 10, **100** | 0.1, 1, **10**, 100 |
| **gamma** | 0.00001,0.00005,0.0001, 0.0005,0.001,0.005,0.01,**0.05** | 0.00001,0.00005,0.0001, 0.0005,0.001,0.005,**0.01**,0.05 | 0.00001,0.00005,0.0001, 0.0005,**0.001**,0.005,0.01,0.05 |

**Table S3. SVM hyperparameter grid search**

| | Gene feature RF | Clinical feature RF | Combined feature RF |
|---|---|---|---|
| **max_features** | 4,**5**,6,7,8,9,10,11,12 | 4,5,6,**7**,8,9,10 | 4,5,6,7,8,**9**,10,11,12 |
| **max_depth** | **4**,5,6,7,8,9,10,11,12 | 4,5,6,**7**,8,9,10 | **4**,5,6,7,8,9,10,11,12 |
| **min_samples_leaf** | **10** | **10** | **10** |
| **n_estimators** | **1000** | **1000** | **1000** |

**Table S4. RF hyperparameter grid search**

| | Gene feature DNN | Clinical feature DNN | Combining part of bimodal DNN |
|---|---|---|---|
| **number of hidden layer** | 2, **3** | 1, **2** | **1** |
| **number of neuron** | 10, **20** | **5**, 10 | 5, **10** |
| **activation function** | **relu**, tanh | relu, **tanh** | relu, **tanh** |
| **learning rate** | 0.002, **0.0002** | **0.002**, 0.0002, 0.00002 | **0.00002** |
| **batch size** | **10** | **10** | **10** |
| **Number of epoch** | **200**, 400, 600, 800 | 200, 400, 600, **800** | epoch_1*: 50, **100** epoch_2**: 0, 50, **100** |
| **optimizer** | **Nadam** | **Nadam** | **Nadam** |
| **l2-regularization term** | **0. 0001** | **0.0001** | **0.0001** |
| **dropout rate** | **0.05** | **0.05** | **0.05** |
| **max_norm constraint** | **1** | **1** | **1** |

**Table S5. DNN hyperparameter grid search**

#### H. Test performance of selected genes as single biomarkers

| | Positive / negative correlation with poor prognosis | AUC | CI |
|---|---|---|---|
| **ELAVL1** | - | 0.5471 | 0.5254 |
| **EGFR** | + | 0.6739 | 0.6170 |
| **BTRC** | - | 0.7258 | 0.6228 |
| **FBXO6** | + | 0.5169 | 0.5195 |
| **SHMT2** | + | 0.6957 | 0.6331 |
| **KRAS** | + | 0.6581 | 0.5894 |
| **SRPK2** | - | 0.7029 | 0.6078 |
| **YWHAQ** | + | 0.6575 | 0.6113 |
| **PDHA1** | + | 0.7032 | 0.6095 |
| **EWSR1** | - | 0.5386 | 0.5083 |
| **ZDHHC17** | - | 0.6507 | 0.5960 |
| **ENO1** | + | 0.6516 | 0.6036 |
| **DBN1** | + | 0.5809 | 0.5566 |
| **PLK1** | + | 0.7125 | **0.6406** |
| **ESR1** | - | **0.7349** | 0.6321 |
| **GSK3B** | + | 0.5836 | 0.5504 |
| **PGR** | - | 0.7062 | 0.6235 |
| **ERBB2** | - | 0.505 | 0.5035 |
| **MKI67** | + | 0.6982 | 0.6166 |
| **PLAU** | + | 0.6132 | 0.5520 |

**Table S6. Test performance of selected genes as single biomarkers**
